# Maternal separation recalibrates prefrontal mitochondrial function and protects against stress-induced negative cognitive bias

**DOI:** 10.64898/2026.07.08.737201

**Authors:** Olivia Stupart, Lucia Marti-Prats, Lorenz MW. Holzner, Sarah Ibegbulam, Amy L. Milton, Rebecca P. Lawson, Andrew J. Murray, Clara Velazquez-Sanchez, Jeffrey W. Dalley

**Affiliations:** Department of Psychology, University of Cambridge, Downing St, Cambridge CB2 3EB, UK; Department of Physiology, Development & Neuroscience, Downing Street, Cambridge, CB2 3EG, UK; Department of Psychiatry, Herschel Smith Building for Brain and Mind Sciences, University of Cambridge CB2 0SZ, UK

**Author notes:** Author for correspondence: Professor Jeffrey W. Dalley, Department of Psychology, University of Cambridge, Downing St, Cambridge CB2 3EB, UK. |. Joint senior authors (Clara Velazquez-Sanchez, Jeffrey W. Dalley).

**Keywords:** Early life stress, Maternal separation, Judgment bias, Ambiguity, Stress resilience, Prefrontal cortex, Mitochondrial function

## Abstract

Ambiguity represents a form of uncertainty in which outcome probabilities cannot be explicitly learned, making decisions dependent on emotional states and cognitive biases. Early-life stress (ELS) increases the risk of adverse mental and physical health outcomes and alters affective processing and learning. ELS may thus affect how ambiguous information is processed, which may depend on interactions with adulthood stress (AS) and mechanistically on bioenergetic mechanisms mediated by top-down cognitive control systems within the prefrontal cortex (PFC). The present study investigated the effects of AS in rats exposed to early maternal separation (MS), a rodent model of ELS, on a task assessing cognitive bias, together with putatively accompanying alterations in PFC mitochondrial function. Cognitive bias was assessed using an ambiguous cue task (ACT) in MS and non-separated control rats tested at baseline and following repeated unpredictable mild stress during adulthood. MS did not affect baseline cognitive bias but increased response latencies. Following AS, control animals showed a significant negative shift in cognitive bias, whereas MS animals were resistant to this shift. MS was also associated with greater PFC mitochondrial respiratory capacity and uncoupling of oxidative phosphorylation following AS. These findings suggest that ELS is associated with a recalibrated phenotype that buffers against the affective consequences of later stress. Enhanced PFC mitochondrial bioenergetics may underlie this resilience, highlighting the importance of developmental context in shaping affective-cognitive responses to stress.

## Introduction

In uncertain situations, the interpretation of ambiguous information is strongly influenced by an individual’s affective state, stress levels and emotional responsiveness^1^. Consequently, a predisposition to make pessimistic interpretations or choices has been widely used as an index of negative affect^2,3^. While transient negative biases may be adaptive under acute stress by facilitating threat detection and promoting cautious responding^3,4^, persistent pessimistic bias is strongly associated with the onset and maintenance of mood and anxiety disorders^5–7^. Early life stress (ELS) is a major risk factor for the development of psychopathology later in life^8–10^. Adversity during sensitive developmental periods induces lasting alterations in emotional processing, stress responsivity, and cognition^11^. However, the consequences of ELS are not uniform and depend on individual patterns of adaptation or maladaptation to subsequent stressors^11–13^. Maternal separation (MS), the most widely used translational model of ELS in rodents, reproduces many behavioural and physiological features of human adversity^14–17^. Importantly, its effects are highly heterogeneous, varying according to procedural parameters^18–20^ and later-life environmental context^21^.

A growing body of evidence suggests that MS induces a latent phenotype, whereby alteration in affective-cognitive processing emerge following subsequent stress exposure, despite the absence of apparent baseline impairments^21,22^. Two conceptual frameworks have been proposed to explain these outcomes. McEwen’s allostatic model^23^ proposes that repeated activation of physiological stress responses leads to cumulative allostatic load, producing persistent changes in stress-response systems, including a dysregulation of the hypothalamic–pituitary–adrenal (HPA) axis and prefrontal–amygdala circuitry. Within this framework, the “multi-hit” or cumulative stress hypothesis posits that ELS sensitises these systems, increasing vulnerability to later stressors, amplifying negative cognitive bias and risk for psychopathology^24,25^. In contrast, the stress inoculation model suggests that moderate early adversity may strengthen adaptive coping mechanisms enhancing resilience to future challenges^26,27^. However, the mismatch hypothesis further highlights an important boundary condition: when later stressors differ in nature or exceed the intensity of prior experiences, these early adaptations may become insufficient or even maladaptive, increasing vulnerability rather than resilience^25^.

Judgement bias depends on neural systems that assign value to uncertain cues and flexibly update behaviour in accordance with an individual’s internal state. Frontal regions, particularly the orbitofrontal (OFC) and prefrontal (PFC) cortices, integrate affective and cognitive information to guide decision-making under ambiguity. These regions encode outcome values, track changing reward contingencies and support behavioural flexibility^28,29^. In rodents, lesions of the OFC alter judgement bias, highlighting the importance of frontal valuations systems in ambiguous cue tasks^30^. Importantly, these valuation processes are highly sensitive to stress. Chronic stress disrupts cognitive processing and promotes negative emotional states^31^, while shifting the interpretation of ambiguous information towards pessimistic responding^4,32^. Such biases are therefore widely used as behavioural markers of affective disorders.

At the cellular level, mitochondria have emerged as key candidates linking stress exposure to cognitive and affective outcomes^33,34^. Beyond ATP production, mitochondria regulate calcium buffering, redox signalling, and apoptotic pathways^35,36^. Stress disrupts mitochondrial function, altering energy metabolism and increasing oxidative stress, leading to a heightened vulnerability to neuronal dysfunction^37,38^, effects that may be particularly important in the energetically demanding PFC region, where efficient mitochondrial function is required to support flexible, goal-directed behaviour^39^. Consistent with this notion, chronic variable stress in mice is associated with the enrichment of mitochondrial gene pathways in the PFC^40^ and similar transcriptional signatures have been reported in individuals with major depressive disorder^41^. While such changes may initially reflect compensatory mechanisms to meet heightened energetic demands, sustained upregulation may have deleterious consequences, ultimately contributing to cellular dysfunction. These findings suggest a mechanistic pathway through which ELS may induce persistent alterations in PFC mitochondrial function capable of influencing neural processes underlying value-based decision-making under ambiguity, especially involving negative affective states.

To investigate this mechanism, the present study assessed whether MS alters the impact of adult stress (AS) on responding to ambiguous cues, and whether behavioural changes are accompanied by alterations in PFC mitochondrial function. MS and control animals were first assessed on a novel auditory ambiguous cue task (ACT) to establish baseline cognitive bias before exposure to a mild inescapable foot-shock procedure designed to induce a negative affective state. Behaviour was subsequently re-assessed, and post-mortem PFC tissue was analysed using high-resolution respirometry and molecular profiling. This experimental design allowed us to test whether MS increases vulnerability to stress-induced pessimistic bias or, alternatively, induces a recalibrated phenotype characterised by altered responsiveness to later stress that is accompanied by corresponding alterations in PFC mitochondrial function.

## Methods

### Subjects

Pregnant Lister-Hooded dams (n = 10; Envigo, Blackthorn, UK) arrived at the vivarium at embryonic day 14 (E14) and remained undisturbed until parturition. During pregnancy and lactation, dams and litters were housed under a 12 h light-dark cycle (lights on 07:00–19:00) with food and water *ad libitum*. At postnatal day (PND) 21, pups (N = 80) were weaned and housed in same-sex groups of four. Food restriction began on PND62, with animals maintained above 85% of predicted free-feeding weight. All procedures were approved by the University of Cambridge Animal Welfare and Ethical Review Body and conducted under UK Home Office Project Licence PP4363352 in accordance with the Animals (Scientific Procedures) Act 1986 (Amendment Regulations 2012) and EU Directive 2010/63/EU.

### Timeline and experimental design

The detailed experimental timeline is shown in **Fig. 1A**. Litters were born between gestational days 22–24 (PND0) and sexed on PND2. After litter adjustment (6–12 pups), animals were allocated to maternal separation (MS; n = 40) or control (n = 40) groups with equal numbers of males and females. From PND62 animals underwent food restriction before training on the ambiguous cue task (ACT). Following training, rats were randomly assigned to AS or control conditions, generating eight experimental groups (sex × MS × AS; n = 10/group). Animals were sacrificed 7 days after the final AS session.

**Figure 1:**
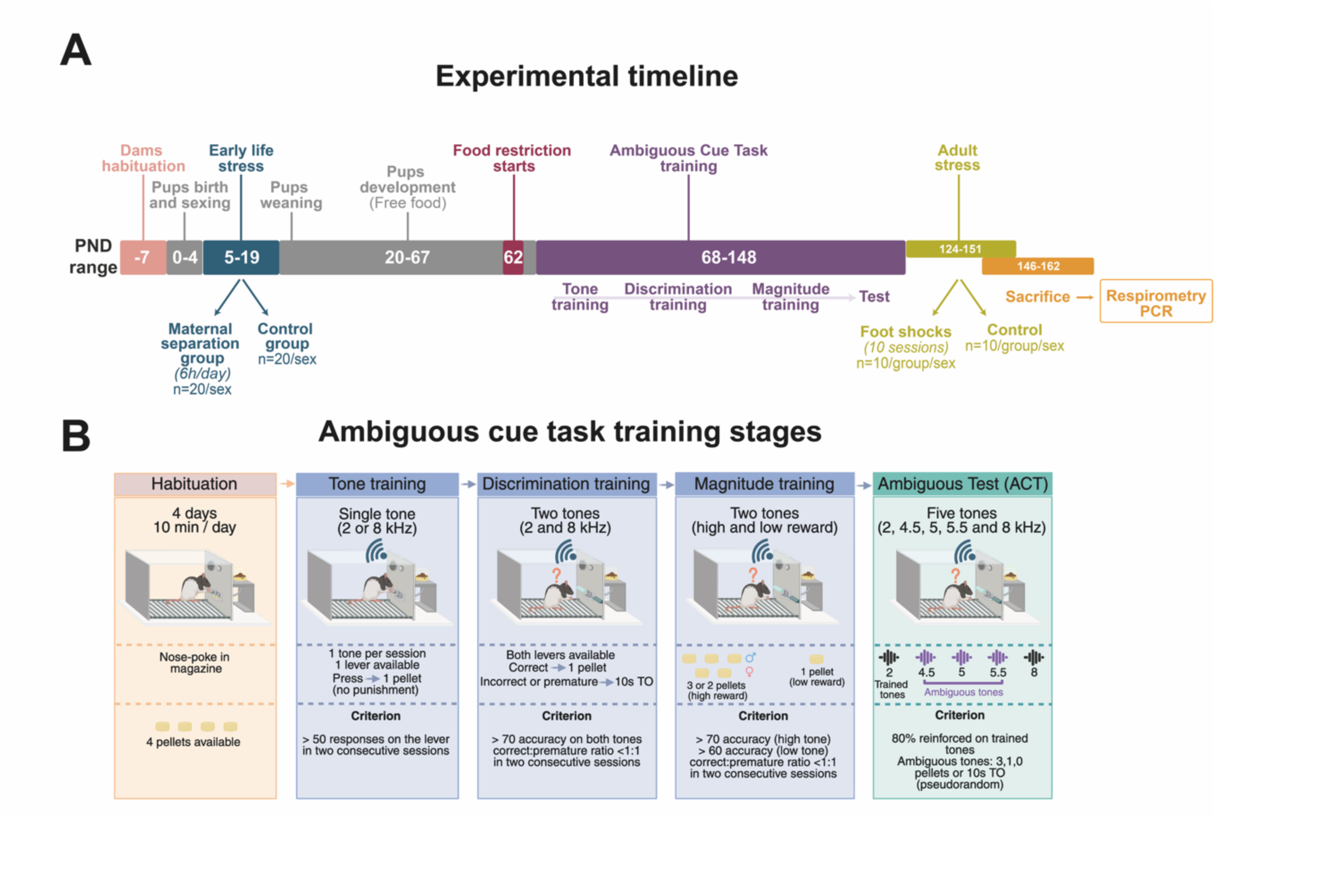
**A) Experimental timeline**. Pregnant Lister Hooded dams arrived at the animal facility on embryonic day (E)14. Upon arrival, dams were left undisturbed with ad libitum access to food and water for at least 7 days before they started littering. Nesting material was provided and litters were delivered via spontaneous labour on E21–22. The day of birth was designated postnatal day (PND)0, and all litters were born within a 24-hour period. Between PND2–4, pups were sexed, and litters were randomly assigned to either maternal separation (MS, n= 20 / sex) or control (n= 20 / sex) conditions. The MS procedure was conducted daily from PND5–19 for 6 hours per day. At PND21, pups were weaned and subsequently housed in groups of four, consisting of animals from mixed litters but matched for sex and experimental condition. Animals were maintained on a 12-hour reverse light cycle with ad libitum access to food and water. Food restriction began at PND62, after which behavioural training on the ambiguous cue task commenced at PND68. In adulthood (PND124), MS and control animals were randomly allocated to either an adult-life stress (AS) group (n= 10 / group / sex) or control (n= 10 / group / sex) condition. The AS protocol consisted of un-cued, unpredictable, and inescapable 10-foot shocks administered over 15 days. At the end of the experiment, animals were sacrificed by decapitation. Brains were rapidly extracted, flash-frozen in isopentane on dry ice, and stored at −80 °C until further analysis. **B) Ambiguous cue task training stages.** Briefly, animals were habituated to the operant chambers (4 days, 10 min/day) before undergoing sequential training stages. During the tone training, a single tone (2 or 8 kHz) was paired with one lever. Pressing the correct lever triggered the delivery of one pellet and no punishment for incorrect responses was programmed. In the discrimination training, both tones were presented pseudo-randomly, requiring the animals to choose the correct lever to obtain the reward. Incorrect or premature responses were punished with a 10 s time out (TO). During magnitude training, one tone was associated with a higher reward (2 pellets for females, 3 for males) and the other with a lower reward (1 pellet). Finally, in the ambiguous test phase, five tones (2, 4.5, 5, 5.5, and 8 kHz) were presented (ratio 4:2:2:2:4), with intermediate tones representing the ambiguous stimuli. Trained tones were reinforced 80% of the trials, while ambiguous tones were pseudo-randomly reinforced (3, 1, or 0 pellets, or TO) to prevent learning of fixed contingencies. Sessions lasted up to 60 min or 140 trials. Figure created in BioRender. https://BioRender.com/qgi3abt

### Maternal separation procedure

MS was carried out as previously described^42^ with minor adaptations. Briefly, from PND5–19, MS litters were separated from the dam for 6 h/day during the dark phase, whereas control litters remained undisturbed except for routine husbandry. During separation, pups were housed together in litter-specific cages within a ventilated incubator (27–28°C) containing bedding, a heat shield, and food and water when developmentally appropriate. Pups were monitored hourly and reunited with the dam following scent transfer using home-cage bedding. Maternal behaviour resumed normally in all litters within 5 min.

### Apparatus: ambiguous cue task

Behavioural experiments were performed in sixteen identical operant chambers located in ventilated and sound-attenuating cubicles (Med Associates, St. Albans, VT). Each chamber was made of aluminium and transparent acrylic plastic with a stainless-steel grid floor (24 x 25.4 x 26.7 cm) and was equipped with a house light (3-W), an illuminated food magazine, a custom-built tone generator, two retractable levels, and an LCD touchscreen on the opposite wall deactivated for this task. Chambers were controlled by Klimbic software (Conclusive Marketing Ltd, UK) and behavioural data were processed using an open-access python script (https://github.com/Jeff-Dalley-Github).

The tone generator consisted of an Arduino-controlled dual piezo-buzzer system capable of delivering 5 different tones (2, 4.5, 5, 5.5 and 8 kHz). Tone frequencies were measured using Advanced Spectrum Analyzer PRO for mobile devices and presented at a sound intensity of approximately-50 and-40 dB.

### Ambiguous cue task (ACT): Training stages

Animals were habituated to the operant boxes (4 days, 10 minutes/day) before starting the training on the ACT. Training sessions lasted for either 60 min or until the maximum trials for that stage were completed, whichever occurs sooner. During Tone Training, animals learned the association between a single tone (2 or 8 kHz) and one lever. Incorrect responses were not punished, and trials were separated by a 5s inter-trial-interval (ITI). During Discrimination Training, both tones were presented pseudo randomly and animals required to respond on the associated lever to obtain reward, while incorrect or premature responses resulted in a 10s timeout (TO) during which the light house was illuminated. During Magnitude Training, one tone predicted a larger reward than the other. Females received two pellets instead of three as the high-value reward to minimise omissions caused by satiation. Once criterion performance was achieved, animals completed the ambiguous test in which the two learnt tones and three novel intermediate tones (4.5, 5 and 5.5 kHz) were presented in a 4:2:2:2:4 ratio. Learnt tones remained reinforced on 80% of trials, whereas ambiguous tones were pseudo-randomly reinforced to prevent stable contingency learning. The details of each training stage are shown in **Fig. 1B**. Cognitive Bias Index (CBI) was calculated as the proportion of high-reward minus low-reward lever responses (range −1 to +1), with higher values indicating a more positive bias.

### Inescapable Adult Stress

Beginning between PND124-151, half of the subjects were exposed to a later-life stressor^21^. The AS procedure was carried out in a context distinct from the ACT using differently configured operant chambers (26 × 26 × 21 cm; Paul Frey Ltd) fitted with a house light kept on throughout the session, and a cotton pad placed beneath the metal grid floor containing three drops of tea tree essential oil. During each 10-min session, animals received 0, 1-or 2-foot shocks (0.5 mA, 0.5 s), delivered pseudo randomly, with no shocks during the first 2 min and no more than 10 shocks per animal across the experiment. Animals underwent 10 stress session over 14 days using an automated Whisker-controlled system^43^.

### Tissue Collection

Seven days after the final stress exposure, rats were briefly anaesthetized with isoflurane (<30 seconds) before decapitation. Brains were rapidly extracted and bisected along the midline.

The PFC from one hemisphere was dissected for mitochondrial respirometry assays while the remain tissue was snap frozen in isopentane at-40°C (Sigma-Aldrich) and stored at-80°C for subsequent molecular analyses.

### High-resolution respirometry

High-resolution respirometry was performed as previously described^44^. Briefly, mitochondrial respiratory function was assessed in PFC using Oxygraph-2k high-resolution respirometers (Oroboros Instruments, Innsbruck, Austria). Following dissection, tissue was maintained in ice-cold BIOPS solution and homogenised in MiR05 at a concentration of 1 mg tissue per 20 µL, using an Eppendorf pestle attached to a DLH overhead stirrer (Velp Scientifica, Usumate, Italy) set at 800 rpm for three 10 sec intervals, with samples cooled on ice between rounds.

Two milligrams of homogenate were added to Oxygraph-2k chambers containing MiR05 at 37°C. Oxygen flux (*J*O_2_) was assessed using a substrate-uncoupler-inhibitor titration (SUIT) protocol as shown in **Table 1**. All assays were performed in duplicate and averaged. All respiration rates were corrected by subtraction of ROX values and normalised either to tissue wet weight or to maximal uncoupled ETS capacity (PGMS*_E_*) to generate flux control ratios (FCRs). In addition, pyruvate-supported OXPHOS coupling efficiency (OCE), an index of mitochondrial respiratory efficiency, was calculated as OCE= (PM*_P_* – PM*_L_*)/PM*_P_*.

**Table 1.** Respiratory control states in high resolution respirometry. Flux Control Ratios (FCR) are calculated by subtracting residual oxygen consumption (ROX) and dividing each respiration rate by the maximal uncoupled ETS capacity (PGMS*_E_*). Table created in BioRender. https://BioRender.com/fd949nn.

| Respiratory state | Substrates used | Concentration (nM) | Corresponding FCR | Pathways probed |
| --- | --- | --- | --- | --- |
| **PM<sub>L</sub>** | Malate<br>Pyruvate<br>ADP | 2<br>10<br>5 | FCR<sub>PML</sub> | Pyruvate supported LEAK |
| **PM<sub>P</sub>** | Cytochrome c | 0.01 | FCR<sub>PML</sub> | Pyruvate supported OXPHOS |
| **PGM<sub>P</sub>** | Glutamate | 20 | FCR<sub>PGML</sub> | Complex I supported OXPHOS |
| **PGMS<sub>P</sub>** | Succinate | 10 | FCR<sub>P/E</sub> | Maximal OXPHOS capacity |
| **OctPGMS<sub>E</sub>** | FCCP | 0.25 μM additions until<br>maximum reached | N/A | Maximal uncoupled ETS<br>capacity. |
| **S<sub>E</sub>** | Rotenone | 0.0005 | FCR<sub>SE</sub> | Complex II supported uncoupled<br>ETS capacity. |
| **ROX** | Antimycin A | 0.0025 | N/A | Non-mitochondrial oxygen<br>consumption. |

### PCR analysis

The contralateral flash frozen PFC was homogenised in Qiazol reagent (Qiagen, Hilden, Germany) and RNA extracted using Quick-RNA Miniprep kit (Zymo Research, Irvine, USA). Reverse transcription was performed using 400ng of RNA (Quantitect Reverse transcription kit, Qiagen). Quantitative PCR was carried out using the Quantinova SYBR green kit (Qiagen) on a LightCycler 480 II (Roche AG, Basel, Switzerland), with 2ng of cDNA used per reaction. Primers are listed in **Table 2**. Gene expression was normalised to the geometric mean of the three most stable housekeeping genes (*GAPDH*, *Eef1*, *ActB*), identified using BestKeeper software^45^, and quantified using the 2-ΔΔCt method^46^.

**Table 2:**
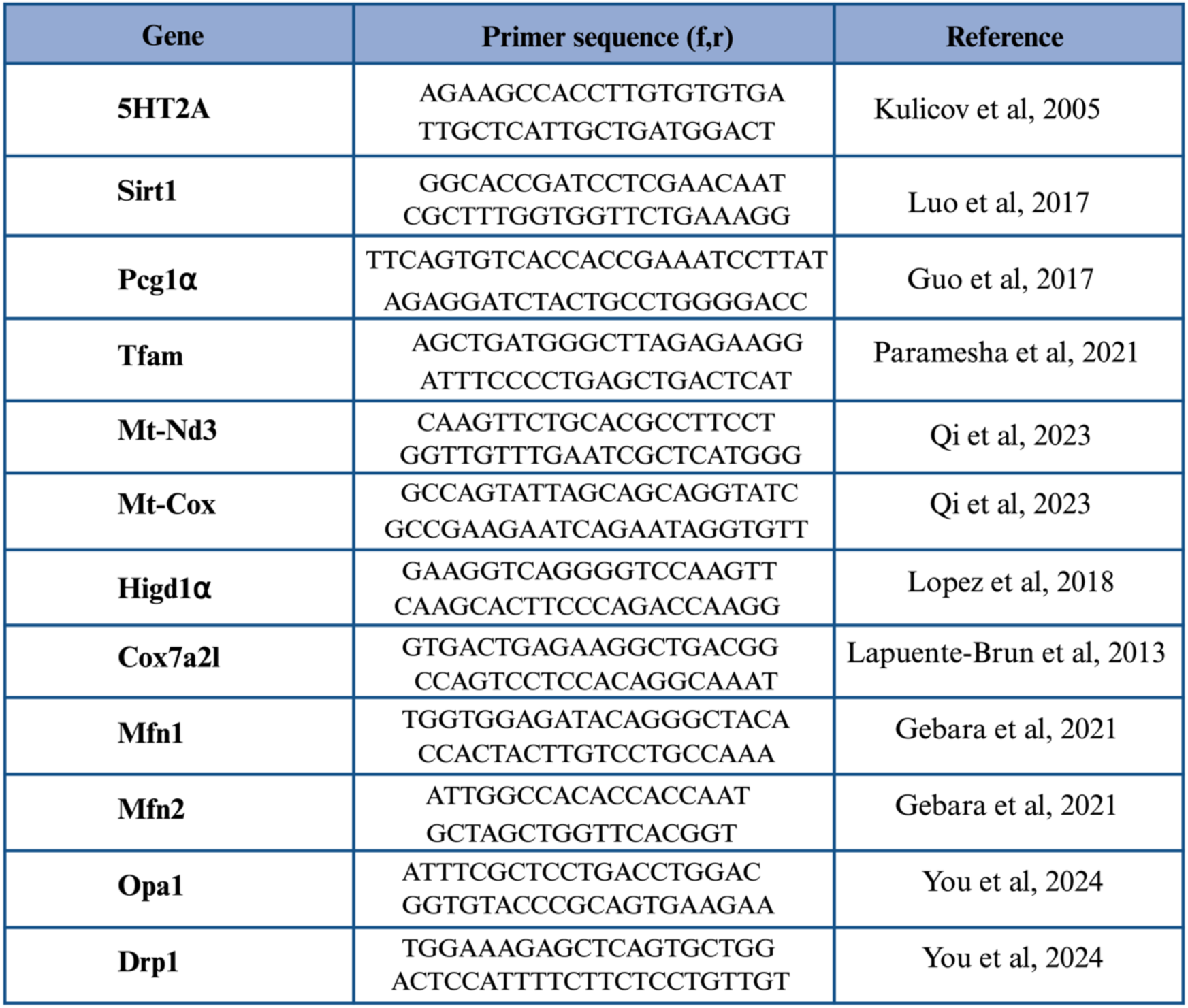
Table of RT-PCR primer sequences. Table created in BioRender. https://BioRender.com/fd949nn. Table references^74,83–90^.

### Data Analysis

Data statistical analysis was carried out using RStudio (version 4.3.2) and figures were generated using BioRender and Adobe Illustrator (Adobe, CA, USA). Data are presented as mean ± SEM or as boxplots (median ± 25% [percentiles] and 5–95% whiskers) with individual data points.

Baseline performance during the Magnitude training was first assessed to ensure stable task acquisition prior testing in the ACT. Behavioural measures (percentage correct, percentage premature, omission and response latency) were analysed using linear mixed-effects models, with stress, sex and session as a fixed factors and relevant combinations of subject, date/time and ACT group as random effects. As females underwent a modified Magnitude Training protocol, sex effects were examined separately.

To investigate the impact of introducing the ambiguous tones during the test phase, the average responses on learnt tones were compared with the averaged responses on ambiguous cues. Data distribution and variance assumptions were assess using Shapiro-Wilk and Levene’s tests, respectively. Group means were analysed using paired t-tests or Wilcoxon signed-rank test with continuity correction when non-parametric alternatives were required. The stability of CBI scores across test sessions was assessed examine using Pearson’s correlation, and CBI was modelled as a function of tone to determine discrimination across ambiguous cues. Model selection was based on likelihood ratio tests (LRTs), Akaike (AIC) and Bayesian (BIC) information criteria. Session number was retained as fixed effect only when it significantly improved model fit. Significant main effects were explored using estimated marginal means (EEMs).

Respirometry and RT qPCR data were analysed using linear models. When significant interaction effects were identified, Tukey’s post hoc comparisons were additionally performed. Significance was set at α ≤ 0.05.

## Results

### Sex dependent differences on the baseline ambiguous cue task

During the five days preceding testing with ambiguous tones, baseline performance remained stable across sessions. A significant main effect of sex was observed for the percentage of correct responses (F(1,223.49) = 32.50, p < 0.0001) and number of omissions (F(1,318.15) = 36.11, p < 0.0001), but not for the percentage of premature responses (data not shown). Controlling for stress and session as fixed effects, and subject and group as random effects, EMMs indicated that males achieved a higher percentage of correct responses (85.9% ± 0.013) compared to females (72.9% ± 0.013; t = 8.00, p < 0.0001) (data not shown). In addition, males made significantly fewer omissions per session than females (t = −10.44, p < 0.0001) (data not shown). Importantly, these sex differences did not indicate a failure of learning in females, since there was no significant sex x session interaction for any measured variable, suggesting that both sexes reached a stable performance over time. Thus, despite differences in response profiles and training adjustments (i.e. reduced reward magnitude), females acquired the task to a comparable level compared with males.

### Maternal separation does not affect cognitive bias

During the ambiguous test phase, previously learnt tones reliably predicted either a high or low reward, whereas the three intermediate sounds, novel to the animals, had no association with reward magnitude. Animals responded more accurately to learnt than ambiguous tones (t = 54.8, p < 0.0001), with a mean difference of 36.18% (95% CI: 34.89–37.48) (**Fig. 2A**), confirming that ambiguous tones were not associated with predictable outcomes. Responses to ambiguous tones were also slower than those to learnt tones (t = 33.17, p < 0.0001) (**Fig. 2B**), with a mean latency of 279 cs and 137 cs, respectively. Omission rated did not differ between trial types (V = 3150.5, p = 0.717), indicating comparable task engagement.

**Figure 2:**
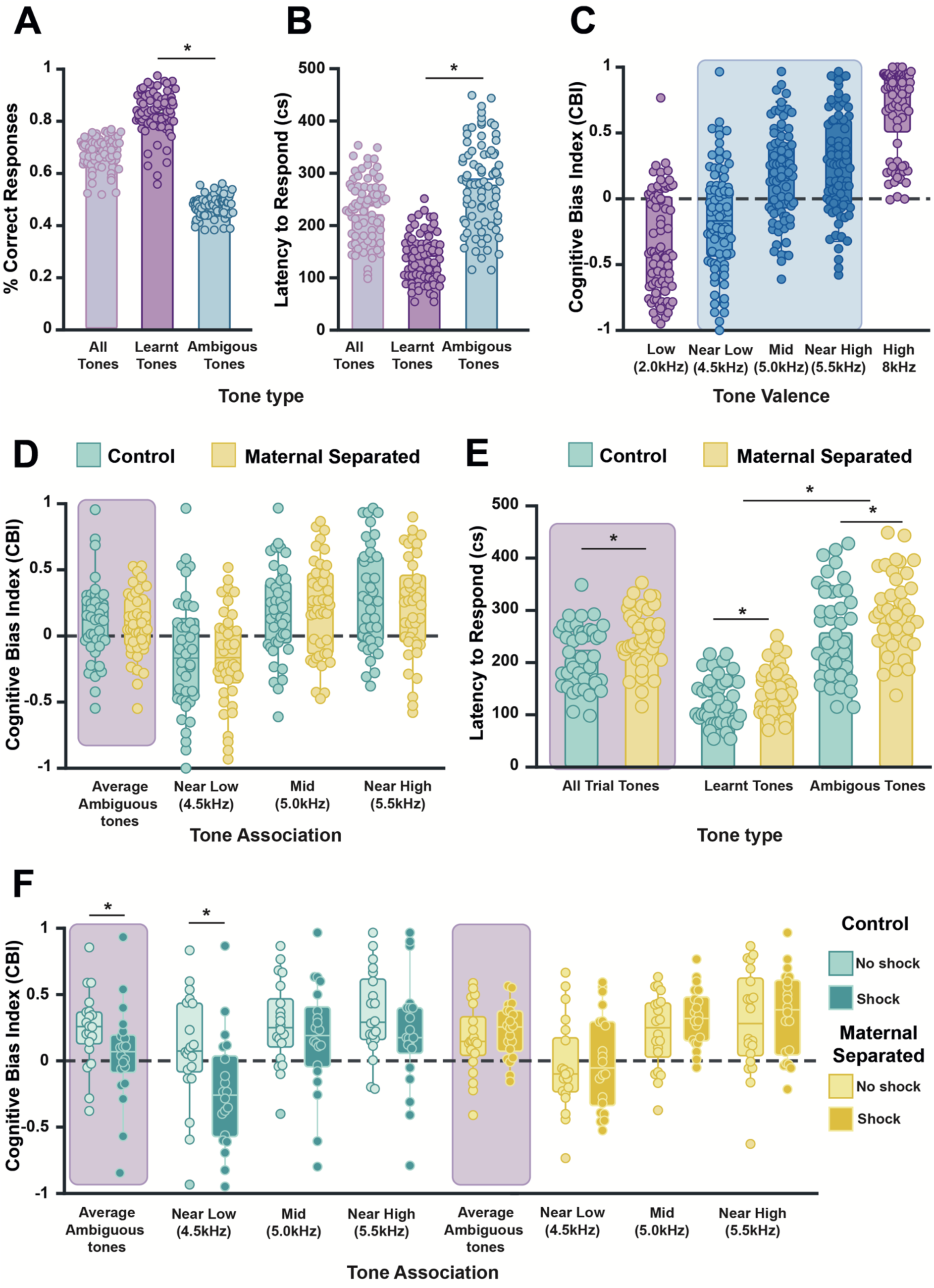
Maternal separation does not modify baseline affective bias. Accuracy on the ambiguous cue task (ACT) measured as the percentage of correct responses was significantly higher for learnt compared to ambiguous tones, indicating that animals reliably recalled reward magnitude associations for familiar tones, while ambiguous sounds remained unpredictable (Fig. 2A). Responses to ambiguous tones were significantly slower than to learnt tones (Fig. 2B). The three ambiguous tones were distributed across the tonal spectrum between the familiar tones and triggered graded cognitive bias responses, corresponding to near-low, middle and near high relative to the learnt tones (Fig. 2C). MS did not significantly affect cognitive bias index (CBI), either averaged across tones or analysed separately for each ambiguous tone (Fig. 2D). In contrast, MS significantly affected response latency, so animals exposed to MS procedure during early life responded slowly overall, as well on both learnt and ambiguous trials (Fig. 2E). **Maternal separation prevents stress-induced negative bias.** Following exposure to AS, animals were re-tested on the ACT. In control animals (non-MS), AS induced a significant negative shift in cognitive bias, whereas this effect was absent in MS subjects, suggesting that previous stress experiences might alter responsiveness to stress exposures later in life (Fig. 2F). Mean CBI scores, averaged across the three ACT sessions, were significantly lower in shocked control animals compared to non-shocked controls (Fig. 2F**, green plots**). No differences were observed between shocked and non-shocked MS animals (Fig. 2F**, yellow plots**). A similar interaction was observed specifically for the near-low ambiguous tone where shocked control animals showed significant lower CBI score values compared to their non-shock controls counterparts, whereas MS animals again remained unaffected by adult shock exposure. * Denotes p < 0.05.

CBI scores were stable across the three pre-AS test sessions (r = 0.635 ± 0.020), indicating that cognitive bias represented a stable individual trait. Incorporating the session variable did not significantly improve model fit. Animals also differentiated between the three ambiguous tones (near-low, middle and near high) (**Fig. 2C**), producing a graded pattern of responding (tone effect: F_(2,620.6)_ = 266.86, p <0.0001). EMMs were −0.2800 ± 0.0351, 0.1430 ± 0.0351, and 0.4100 ± 0.0352 for the near-low, middle and near-high tones, respectively, with all pairwise comparisons significant (p <.0001).

MS did not alter baseline cognitive bias. No differences were observed in percentage accuracy, CBI or omission rates, either overall or when learnt and ambiguous trials were analysed separately (**Fig. 2D**). However, MS significantly increased response latency across trials (F_(1,76)_ = 10.9, p = 0.0015), including learnt (F_(1,73)_ = 7.9, p = 0.0062) and ambiguous trials (F_(1,76)_ = 6.84, p = 0.0107) (**Fig. 2E**). Post-hoc comparisons confirmed longer response latencies in MS animals across all trial types (all trials: t = 3.30, p = 0.0015; learnt trials: t = 2.82, p = 0.0062; ambiguous trials: t = 2.62, p = 0.011).

### Unpredictable adult stress affects affective bias in control but not maternal separated animals

To determine whether AS induced a pessimistic expectation bias, animals were exposed to an inescapable and unpredictable mild foot-shock and subsequently re-tested on the ACT. In animals without prior ELS, AS induced a significant negative shift in cognitive bias. In contrast, this effect was absent in MS animals, suggesting that ELS altered responsiveness to later stress exposure. Analysis of mean CBI across all ambiguous tones revealed a significant MS x AS interaction (F_(1,69.5)_ = 6.24, p = 0.015). This interaction was driven by a significant reduction in CBI in control animals following AS, with EMMs decreasing from 0.2259 ± 0.0071 in non-shocked controls to 0.0413 ± 0.0071 following shock exposure (t = −2.50, p = 0.015) (**Fig. 2F, green plots**). Notably, no significant effect of AS was observed in MS animals. CBI values remained comparable between shocked and non-shocked MS groups (non-shock: EMM = 0.1601 ± 0.0071; shock: EMM = 0.2365 ± 0.0071; t = 1.037, p = 0.303) (**Fig. 2F, yellow plots**). A similar interaction was observed specifically for the near-low ambiguous tone (F_(1,68.82)_ = 7.65, p = 0.007). Control animals exposed to AS showed a significant reduction in CBI compared to non-shocked controls (t = −3.114, p = 0.0027), whereas no significant differences were detected between shocked and non-shocked MS animals (t = 0.802, p = 0.425) (**Fig. 2F)**.

AS did not significantly affect the percentage of correct responses during task performance, either overall or when the ambiguous tones were analysed separately. Similarly, shock exposure did not alter response latency to learnt tones. However, MS animals continued to exhibit generally slower response latencies across conditions (F_(1,72)_ = 66.35, p < 0.0001). For ambiguous tone trials, shock exposure was associated with a reduction in response latency in females but not in males; however, this effect was not significant following *post-hoc* testing. Shock exposure also had no significant effect on number of omitted trials, irrespective of whether omissions occurred during learnt or ambiguous tone presentations (data not shown).

### Maternal separation increases prefrontal OXPHOS capacity and increases adult stress-induced mitochondrial leak

To investigate the effect of MS and its interaction with AS, high-resolution respirometry was performed on PFC tissue collected from the same animals that underwent ACT testing. MS was associated with greater maximal OXPHOS capacity (PGMS*_P_*: F_(1,71)_ = 4.24, p = 0.043), and maximal electron transfer system capacity (PGMS*_E_*: F(1,71) = 4.81, p = 0.032) compared with control animals. No differences were observed in Complex I-or Complex II-linked respiration. However, a significant MS x AS interaction was observed for Complex I-linked LEAK respiration (PM*_L_*: F_(1,71)_ = 4.78, p = 0.032). Post-hoc analysis showed that PM*_L_* respiration was increased in MS animals following AS (p = 0.044), whereas no differences were observed between control groups (**Fig. 3A)**. Consistent with this finding, the LEAK-associated flux control ratio showed a similar pattern (**Fig. 3B)**, while no other flux control ratios were altered, suggesting that our findings reflect broad alterations in mitochondrial organisation or abundance, rather than changes at the level of specific ETS complexes.

**Figure 3:**
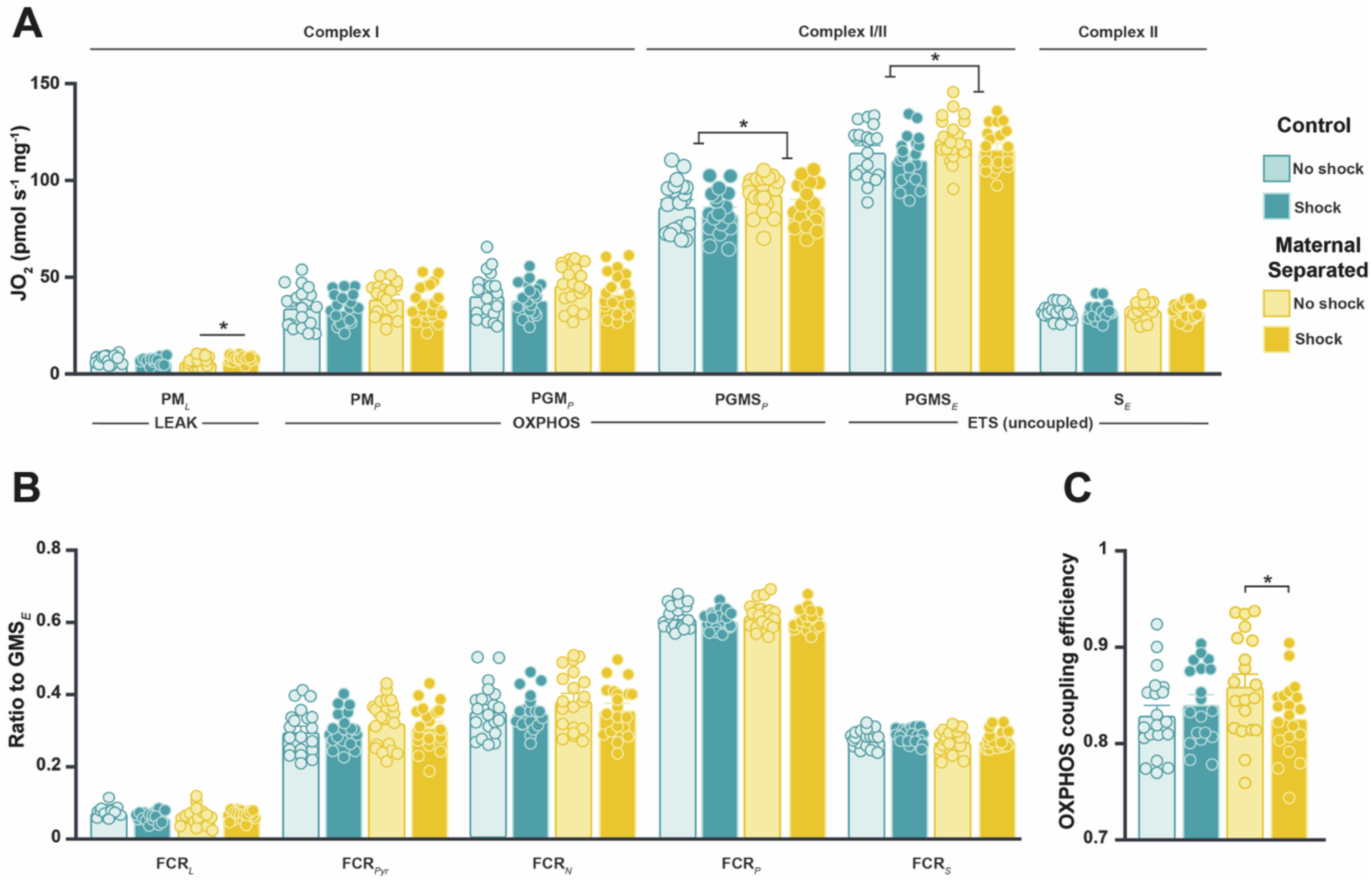
**Effect of maternal separation and adult stress on mitochondrial respiratory function in the prefrontal cortex**. Mitochondrial respiratory capacity was assessed in prefrontal cortex homogenates from either control or MS animals exposed or not to AS using high resolution respirometry. Oxygen consumption was measured after sequential addition of mitochondrial substrates, uncouplers and inhibitors to determine wet mass adjusted respiratory states, including LEAK respiration (PM*_L_*), OXPHOS capacity (PM*_P_*, PGM*_P_,* PGMS*_P_*), maximal electron transport system capacity (PGMS*_E_*, S*_E_*). MS was associated with significantly greater maximal OXPHOS (PGMS*_P_*) and maximal electron transfer system capacity (PGMS*_E_*) compared to control animal. Moreover, MS non-shock animals showed reduced LEAK respiration (PM*_L_*) compared to their MS shock counterparts. In contrast, adult shock exposure was associated with a trend towards reduced maximal OXPHOS capacity (Fig. 3A). Flux control ratios (FCRs) were additionally calculated and rates normalised to PGMS*_E_*. FCR*_L_* = PM*_L_*/PGMS*_E_*; FCR_Pyr_ = PM*_P_*/PGMS*_E_*; FCR*_N_* = PGM*_P_*/PGMS*_E_*; FCR*_P_* = PGMS*_P_*/PGMS*_E_*; FCRS = S*_E_*/PGMS*_E_* (Fig. 3B). OXPHOS coupling efficiency (OCE = (PM*_P_* - PM*_L_*)/PM*_P_*), an index of mitochondrial respiratory efficiency was also calculated. Consistent with the increase in LEAK respiration, OCE was significantly reduced in maternally separated animals exposed to AS compared to MS animals that were not exposed to AS, indicating reduced mitochondrial coupling efficiency. In addition, coupling efficiency was greater in non-shock MS animals relative to stressed MS animals (Fig. 3C). * Denotes p < 0.05

Given the alterations in LEAK respiration, OXPHOS coupling efficiency (OCE; **Fig. 3C**) was calculated. A significant MS x AS interaction was revealed (F_(1,71)_ = 4.34, p = 0.041), with OCE significantly reduced in stressed MS animals compared with non-stressed MS (p = 0.032), indicating reduced mitochondrial coupling efficiency. These findings indicate that MS was associated with increased PFC mitochondrial respiratory capacity, whereas AS selectively increased LEAK respiration and reduced coupling efficiency in MS animals.

### Maternal separation alters mitochondrial gene expression in the prefrontal cortex

To investigate molecular substrates of the altered mitochondrial respiration phenotype, RT-qPCR was performed on PFC samples (**Fig. 4)**. MS did not affect expression of the serotonin *5HT2A* receptor, which has previously been shown to regulate neuronal mitochondrial function^47,48^, although AS was associated with a slight reduction of 5HT2A expression (F_(1,61)_ = 4.02, p = 0.050) (**Fig. 4A)**. We therefore examined the downstream signalling cascade genes associated with *5HT2A* involved in mediating mitochondrial function, *Sirt1* and *Pgc1α.* A significant AS x sex interaction was found on *Sirt1* expression (F_(1,60)_ = 5.26, p = 0.025), where *post-hoc* tests indicated an effect of AS only in males (p = 0.006), whereas *Pgc1α* was unchanged (**Fig. 4B)**.

**Figure 4:**
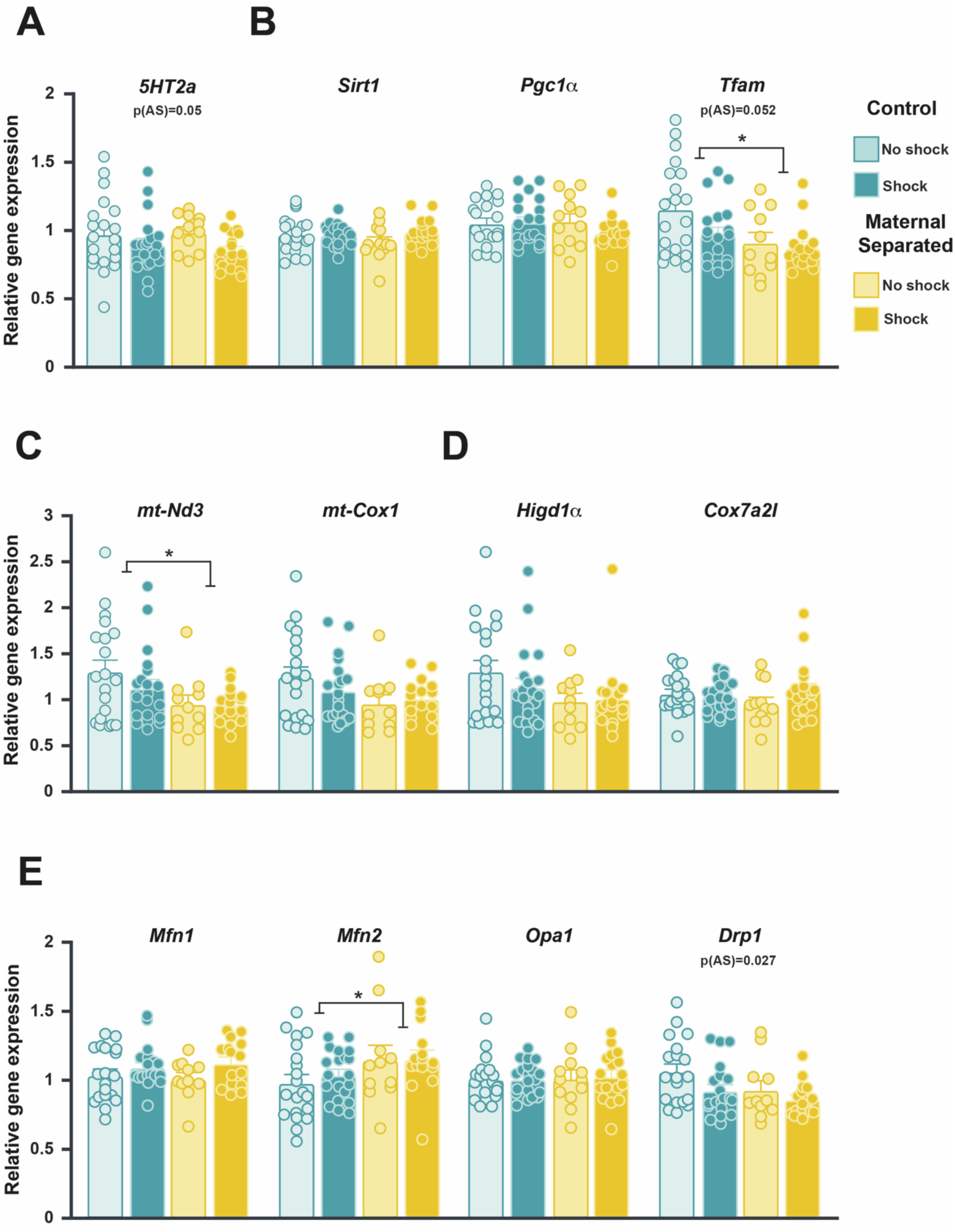
Expression of mitochondria related genes in the prefrontal cortex of rats exposed to maternal separation and/or adult stress. The contralateral prefrontal cortex of the same either control or MS animals exposed or not to AS was used to measure gene expression by RT-qPCR. Data were normalized to the geometric mean of *GADPH*, *Eef1* and *Actb* expression using the 2^-ΔΔCt^ method. Expression of HT2A receptor (Fig. 4A), regulators of mitochondrial activity and gene expression: *Sirt1*, *Pgc1a* and *Tfam* (Fig. 4B), mitochondrial respiratory complex components *mt-Nd3* and *mt-Cox1* (Fig. 4C), the supercomplex assembly factors *Higd1a* and *Cox7a2l* (Fig. 4D) and regulators of mitochondrial fusion and fission, *Mfn1, Mfn2, Opa1* and *Drp1*(Fig. 4E) was analysed. PCR data indicated that AS exposure was associated with reduced expression of *5HT2A* receptor and *Drp1*, together with a near significant reduction in *Tfam* expression. However, the expression of the upstream regulators *Sirt1* and *Pgc1a* and downstream mitochondrial targets *mt-Nd3* and *mt-Cox1* remained unaffected. Contrary, MS was associated with a decreased expression of *Tfam* and mitochondrially encoded respiratory genes *mt-Nd3* and *mt-Cox1*, as well as the fission mediator *Drp1* and with an increased expression of the fusion-related gene *Mfn2.* These findings are consistent with a potential shift towards enhanced mitochondrial fusion following early life stress.

MS significantly reduced expression of *Tfam* (F_(1,58)_ = 6.75, p = 0.012) (**Fig. 4B)**, a key downstream target of the master metabolic regulator *Pgc1α,* and the mitochondrially encoded complex I subunit *mt-Nd3* (F_(1,58)_ = 6.46, p = 0.014) (**Fig. 4C**). However, no significant effects were observed for the *mt-Cox* gene, the main subunit of the cytochrome c oxidase i.e Complex IV, *Higd1α,* a key survival factor during periods of metabolic stress, such as low oxygen or low sucrose levels, or *Cox7a2l*, a structural component involved in respiratory supercomplex scaffolding (**Figs. 4C-D**). To assess mitochondrial dynamics, expression of key genes involved in mitochondrial fusion and fissions was also measured. MS significantly increased the expression of the fusion protein *Mfn2* (F_(1,58)_ = 4.98, p = 0.030), whereas *Mfn1* and *Opa1* were unchanged (**Fig. 4E)**. In contrast, AS significantly reduced expression of the fission regulator *Drp1* (F_(1,58)_ = 5.18, p = 0.027), (**Fig. 4E)**.

Overall, MS and AS were associated with different transcriptional profiles. AS was associated with reduced *5HT2A* and *Drp1* expression, whereas MS was associated with reduced *Tfam* and *mt-Nd3* expression together with increased *Mfn2,* consistent with altered mitochondrial dynamics following ELS.

## Discussion

The present study investigated whether MS alters affective bias in adulthood and whether an interaction exists with later life stress that affects PFC mitochondrial function. We specifically hypothesised that additional stress in MS animals may interfere with the process of top-down cognitive control of stress mediated by the PFC. We report that MS did not alter baseline affective bias, measured using the ACT, although MS animals exhibited longer response latencies and were resistant to the stress-induced negative shift in affective bias observed in control animals. This behavioural profile was accompanied by increased mitochondrial respiratory capacity within the PFC and altered mitochondrial coupling following AS. These findings suggest that ELS may induce persistent alterations in both affective-cognitive processing and PFC bioenergetics, although the relationship between these phenomena remains to be established.

Ambiguous cue tasks are widely used in clinical and preclinical research to assess affective bias and the interpretation of uncertainty under emotional challenge^2,3,22,49,50^. Consistent with previous findings^22^, both control and MS animals exhibit a positive baseline cognitive bias, indicating that MS did not induce a persistent negative affective phenotype under neutral testing conditions. This finding is in line with previous rodent studies showing that similar MS procedures produced little or no overt behavioural outcomes when assessed in adulthood^51,52^, and with human studies in which healthy and depressed individuals often perform similarly when evaluating clearly positive, negative or neutral stimuli, showing comparable levels of accuracy and response latencies^50,53,54^. However, when the stimuli became ambiguous, differences between the groups emerged with depressed individuals being more likely to interpret ambiguous emotional cues as negative^50^.

Ambiguity is a specific form of uncertainty in which outcomes probabilities cannot be explicitly estimated, requiring decisions to rely more on internal estates and prior expectations^55,56^. These processes engage PFC cortical networks involved in uncertainty and behavioural flexibility^29,57^. Consistent with this framework, ambiguity per se was not sufficient to distinguish MS from control animals, whereas behavioural discrepancies emerged after the addition of AS. This suggests that MS altered the response to subsequent challenges rather than their baseline interpretation of ambiguous cues. One possible explanation for this latent phenotype is that the long-term neurobehavioral consequences of MS depend on how ELS shapes the maturation of the hypothalamic-pituitary-adrenal (HPA) axis^58,59^. Although alterations in HPA function may not produce overt behavioural abnormalities under baseline test conditions, these may modify responses to later stressors resulting in differential behavioural, physiological and neuroendocrine outcomes across the lifespan^60,61^. Supporting this interpretation, Shi *et. al*. reported that behavioural differences following predictable MS stress only become apparent after subsequent restraint stress exposure, whereas no baseline differences were observed between MS and control animals^62^. Our findings are therefore consistent with the mismatch hypothesis, which proposes that predictable ELS does not necessarily increase vulnerability to later stress but may recalibrate stress response systems and protect against later stressor exposures^63^. Indeed, findings in humans and non-human primates suggest that moderate or predictable ELS can recalibrate stress-response systems, resulting in reduced physiological responses to later stress and improved coping^27,64–68^.

Although MS did not alter baseline cognitive bias, MS animals exhibited longer response latencies than controls. Response latencies may capture subtle alterations in the cognitive and affective processes underlying decision-making that are not evident from choice behaviour alone. Supporting this interpretation, increased anxiety in humans is associated with slower responding despite preserved task performance^69^, suggesting a reduced processing efficiency rather than impaired decision accuracy. Likewise, early life stress has been associated with reduced processing speed and executive function in humans^70^ and similarly, we previously observed that rodents exposed to MS exhibit slower decision latencies despite comparable task performance^21^. Thus, the prolonged response latencies observed in MS animals may reflect altered processing of ambiguous information or greater deliberation under uncertain trials, without necessarily affecting the overall accuracy of performance.

Recent research implicates mitochondrial function in shaping behavioural responses to stress, with early life adversity inducing persistent changes in mitochondrial biology, that might influence vulnerability or resilience to subsequent challenges^71–73^. Consistent with this literature, MS animals exhibited greater mitochondrial respiratory capacity within the PFC than controls, irrespective of AS exposure. Although the functional significance of this phenotype remains uncertain, enhanced mitochondrial respiratory capacity has been associated with adaptive behavioural responses to stress^33,34,74^. For example, Hollis *et al* demonstrated that low anxious rats, exhibited greater complex I and II protein levels, mitochondrial respiration and ATP production than high anxious animals. While pharmacological inhibition of these complexes increased susceptibility to social subordination, activation protected vulnerable animals from stress-induced subordinate status. These findings tentatively suggest that enhanced mitochondrial respiratory capacity leads to adaptive responses to stress, although whether it contributes directly to behavioural resilience remains unclear. Because the brain relies predominantly on oxidative phosphorylation to meet its substantial energetic demands^75^, the increased respiratory capacity observed in MS animals might reflect persistent alterations in PFC bioenergetic regulation established by early life experience. We hypothesise that such adaptations may increase the energetic reserve available during stress and could contribute to maintaining flexible valuation and decision-making processes, consistent with the preserved cognitive bias observed in MS animals following AS exposure.

To investigate molecular correlates of the altered mitochondrial respiratory phenotype, we investigated the expression of genes involved in mitochondrial biogenesis, respiratory function and mitochondrial fission/fusion dynamics within the contralateral PFC. Overall, MS and AS were associated with different transcriptional profiles. Whereas adults stress primarily altered genes involved in serotonergic signalling and mitochondrial regulation, MS was associated with altered expression of genes related to mitochondrial transcription and dynamics. Specifically, MS animals showed reduced expression of *Tfam*, together with lower expression of the mitochondrially encoded respiratory genes *mt-Nd3* and *mt-Cox1* alongside increased *Mfn2* expression. This profile contrasts with previous studies showing increased expression of *mt-Nd3* and *mt-Cox1* following chronic restraint stress in the PFC^40^, suggesting that early-and later life stress may engage mitochondrial pathways differently. Given the central role of *Tfam* in regulating mitochondrial DNA transcription and replication^76^, the coexistence of reduced mitochondrial transcript expression and enhanced respiratory capacity argues against a simple increase in mitochondrial content. However, transcript abundance does not necessarily predict protein expression, as post-transcriptional regulation, translational efficiency and protein turnover influence the relationship between mRNA and protein levels^77,78^. Consequently, these findings are more consistent with qualitative changes in mitochondrial organisation than increased mitochondrial content. Supporting this idea, MS was also associated with increased expression of the mitochondrial fusion regulator *Mfn2* and reduced expression of the fission mediator *Drp1*. Although this molecular profile has not previously been investigated following MS, increased *Mfn2* expression has been associated with enhanced mitochondrial function and reduced anxiety-and depression-like behaviours^74^. Likewise, Gebara *et al* found that *Mfn2* overexpression restored both mitochondrial impairments and behavioural outcomes in stress-vulnerable animals. These findings raise the possibility that MS is associated with a shift towards a more fused mitochondrial network, a state linked to enhanced mitochondrial connectivity, oxidative metabolism and adaptation to metabolic stress^79–81^. While this interpretation requires confirmation at the protein and morphological levels, our findings are compatible with a persistent reorganisation of PFC mitochondrial architecture accompanying the enhanced respiratory capacity.

AS selectively reduced OXPHOS coupling efficiency in MS rats compared with non-stressed controls. Although reduced coupling efficiency is often interpreted as diminished energetic efficiency, mild mitochondrial uncoupling has also been proposed as an adaptive mechanism that limits oxidative stress and calcium overload while promoting mitochondrial biogenesis and turnover^82^. In the present study, reduced coupling efficiency occurred alongside greater respiratory capacity and preservation of cognitive bias following AS, suggesting that the mitochondrial adaptations observed in MS animals are more consistent with altered bioenergetic regulation than overt mitochondrial dysfunction. Importantly, the present results do not establish that these mitochondrial changes underlie this behavioural phenotype. Rather, resilience to stress-induced negative shift in cognitive bias was accompanied by enhanced respiratory capacity and altered coupling efficiency. One possibility is that the larger respiratory capacity observed in MS animals offered sufficient energetic reserve to compensate the reduction in coupling efficiency and maintain energetic homeostasis. These findings suggest that ELS is associated with persistent alterations in mitochondrial energy regulation that accompany the behavioural response to later stress.

In summary, MS was associated with resilience to the affective cost of AS experiences. Animals exposed to ELS were protected from the stress-pessimistic bias observed in controls and displayed a different mitochondrial phenotype characterised by greater respiratory capacity and altered mitochondrial regulation. While these bioenergetic adaptations may contribute to the observed resilience phenotype, their long-term implications remain unclear. Although they may initially represent compensatory mechanisms that support adaptation to stress, prolonged upregulation of mitochondrial activity could also promote cellular stress and maladaptive neural function. Future studies examining oxidative stress, mitochondrial integrity and behavioural outcomes will be important to establish whether these adaptations represent a stable correlate of resilience or a bioenergetic states imbalance that confers vulnerability to stress later in life.

## Acknowledgements.

We are grateful to Amy Milton as the Home Office project licence holder. We acknowledge the support of Isaac Newton Trust – School of Biological Sciences Interdisciplinary Seed Funding (University of Cambridge) awarded to JWD and AJM.

## Author contributions

OS, CVS and JWD together designed the study; CVS, LMP and LMWH conducted the measurements; OS and CVS performed data analyses; OS and CVS wrote the first draft of the manuscript, which was edited by JWD, LMP, LMWH, SI, ALM, RPL and AJM; CVS and JWD drafted the revised manuscript; all authors approved the initially submitted and revised manuscripts.

## Data availability

Data are available upon reasonable request from the corresponding author. For the purpose of open access, the authors have applied a Creative Commons Attribution (CCBY) licence to any Author Accepted Manuscript version arising from this submission.

## Ethical approval

Experiments were conducted in accordance with the Animals (Scientific procedures) Act, 1983, Amendment Regulations (2012) on project licence PP4363352 following ethical review by the University of Cambridge Animal Welfare and Ethical Review Body (AWERB).

